# An E460D substitution in the NS5 protein of tick-borne encephalitis virus confers resistance to the inhibitor Galidesivir (BCX4430) and also attenuates the virus for mice

**DOI:** 10.1101/563544

**Authors:** Ludek Eyer, Antoine Nougairède, Marie Uhlířová, Jean-Sélim Driouich, Darina Zouharová, James J. Valdés, Jan Haviernik, Ernest A. Gould, Erik De Clercq, Xavier de Lamballerie, Daniel Ruzek

## Abstract

The adenosine analogue Galidesivir (BCX4430), a broad-spectrum RNA virus inhibitor, has entered a Phase 1 clinical safety and pharmacokinetics study in healthy subjects and is under clinical development for treatment of Ebola virus infection. Moreover, Galidesivir also inhibits the reproduction of tick-borne encephalitis virus (TBEV) and numerous other medically important flaviviruses. Until now, studies of this antiviral agent have not yielded resistant viruses. Here, we demonstrate that an E460D substitution, in the active site of TBEV RNA-dependent-RNA-polymerase (RdRp), confers resistance to Galidesivir in cell culture. Stochastic molecular simulations indicate that the steric freedom caused by the E460D substitution increases close electrostatic interactions between the inhibitor and the interrogation residue of the TBEV RdRp motif F, resulting in rejection of the analogue as an incorrect/modified nucleotide. Galidesivir-resistant TBEV exhibited no cross-resistance to structurally different antiviral nucleoside analogues, such as 7-deaza-2’-*C*-methyladenosine, 2’-*C*-methyladenosine and 4’-azido-aracytidine. Although, the E460D substitution led only to a subtle decrease in viral fitness in cell culture, Galidesivir-resistant TBEV was highly attenuated *in vivo*, with 100% survival rate and no clinical signs observed in infected mice. Our results contribute to understanding the molecular basis of Galidesivir antiviral activity, flavivirus resistance to nucleoside inhibitors and the potential contribution of viral RdRp to flavivirus neurovirulence.

**Importance:** Tick-borne encephalitis virus (TBEV) is a pathogen that causes severe human neuroinfections in large areas of Europe and Asia and for which there is currently no specific therapy. We have previously found that Galidesivir (BCX4430), a broad-spectrum RNA virus inhibitor, which is under clinical development for treatment of Ebola virus infection, has a strong antiviral effect against TBEV. For any antiviral drug, it is important to generate drug-resistant mutants to understand how the drug works. Here, we produced TBEV mutants resistant to Galidesivir and found that the resistance is caused by a single amino acid substitution in an active site of the viral RNA-dependent RNA polymerase, an enzyme which is crucial for replication of viral RNA genome. Although, this substitution led only to a subtle decrease in viral fitness in cell culture, Galidesivir-resistant TBEV was highly attenuated in a mouse model. Our results contribute to understanding the molecular basis of Galidesivir antiviral activity.

## Introduction

Flaviviruses (family *Flaviviridae*, genus *Flavivirus*) cause widespread human morbidity and mortality throughout the world. These viruses are typically transmitted to humans by mosquito or tick vectors. Tick-borne encephalitis virus (TBEV) is a typical flavivirus transmitted by *Ixodes* spp. ticks. TBEV is a causative agent of tick-borne encephalitis (TBE), a severe and potentially lethal neuroinfection in humans (Baier, 2011). The disease is prevalent in the sylvatic areas of Europe and Asia with more than 13,000 cases of TBE being reported annually (Dumpis, Crook, and Oksi 1999; Heinz and Mandl 1993). Clinical presentation of TBE ranges from mild fever to severe encephalitis or encephalomyelitis. In many cases, survivors of TBE suffer long-term or even permanent debilitating sequelae (Ruzek, Dobler, and Mantke 2010). As for other flaviviral infections, there is no specific treatment for TBE, other than supportive therapy. Thus, the search for antiviral agents for specific chemotherapy of TBE and relative viruses is urgent.

Among the different strategies aimed at inhibiting virus or cell components involved in TBEV replication, the viral nonstructural NS5 protein, an RNA-dependent RNA-polymerase (RdRp), has become an attractive target for specific and effective inhibition of viral replication, with limited measurable effect on the host cells (Eyer et al., 2018). Several molecules, mainly nucleoside analogues, were found to be potent inhibitors of the TBEV NS5 polymerase activity (Eyer et al., 2015; 2016; 2017a,b; 2018). The mechanism of action of these nucleoside analogues is based on their initial metabolization to the active triphosphate (nucleotide) form by cellular kinases and subsequent incorporation into the nascent genome by the RdRp, leading to premature chain termination (Eyer et al., 2018). Galidesivir (also known as BCX4430 or Immucillin-A, Figure 1A) is an adenosine analogue with two structural modifications: (i) it is a C-nucleoside characterized by a C-glycosidic bond instead of the usual N-glycosidic bond, and (ii) the furanose oxygen has been replaced by an imino group (Warren et al., 2014). Galidesvir is known to have a broad-spectrum antiviral effect against more than 20 different medically important RNA viruses across nine different virus families (flaviviruses, togaviruses, bunyaviruses, arenaviruses, paramyxoviruses, coronaviruses, filoviruses, orthomyxoviruses and picornaviruses) (Warren et al., 2014; Westover et al., 2018; Julander et al., 2014, 2017; Taylor et al., 2016; De Clercq, 2016). Low micromolar levels of Galidesivir were previously shown to inhibit TBEV with no or negligible cytopathic effect on the host cells (Eyer et al., 2017a). A Phase 1 clinical safety and pharmacokinetics study in healthy subjects has been completed, and at present, Galidesivir is under clinical development as an antiviral drug for treatment of Ebola virus infection (Taylor et al., 2016). The broad-spectrum antiviral activity makes this drug a promising candidate for development of therapy not only for Ebola virus infection but also other important diseases caused by various RNA viruses, including TBEV.

**Fig 1.**
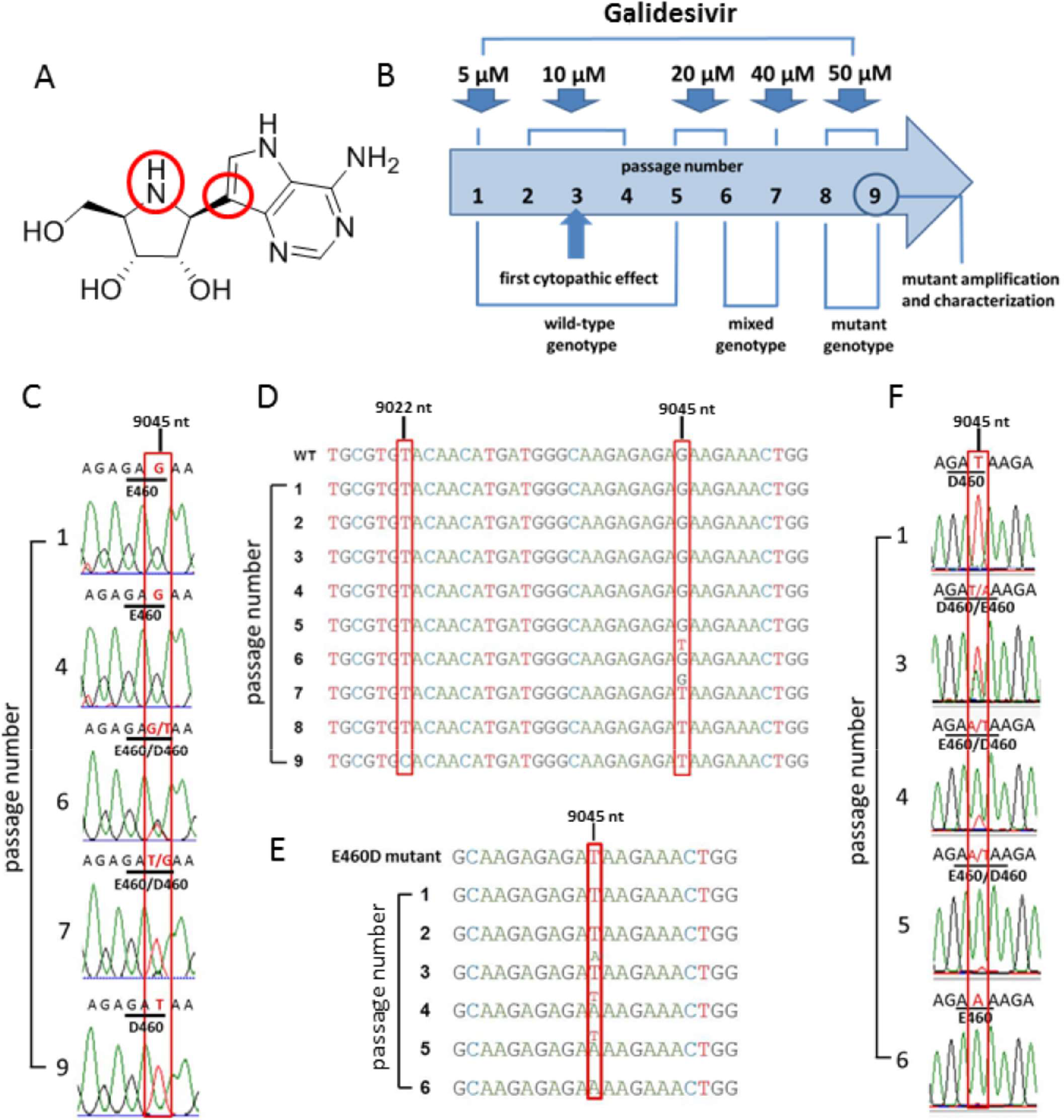
TBEV resistance to Galidesivir is associated with a single mutation in the NS5 gene. (A) The structure of the nucleoside analogue Galidesivir. (B) Scheme of the selection process for generation of TBEV resistance to Galidesivir. TBEV was serially passaged in PS cells in the presence of increasing Galidesivir concentrations. (C) Whole-genome sequence analysis of the passaged viruses revealed a mutation at the amino acid position 460 in the NS5 protein, changing the glutamic acid residue to aspartic acid. (D) In the E460D/Y453H mutant, both amino acid changes E460D (nucleotide position G9045T) and Y453H (nucleotide position and T9022C) were detected in the NS5 protein after 5 and 8 passages in the presence of Galidesivir, respectively. (E) Rapid re-induction of the wild-type genotype in PS cells following serial passage of TBEV in PS cells in the absence of Galidesivir. (F) During serial passage of the E460D mutant virus in PS cells in the absence of galidesivir, the amino acid codon GAT (in the mutant) was changed to the codon GAA (in the revertant).

Although Galidesivir has been studied intensively and is known to inhibit a wide range of RNA viruses, there are no published reports of resistance to this compound. Experience with the treatment of other RNA virus infections shows, that resistance can develop rapidly with any of the direct-acting antiviral agents (Poveda et al., 2014; Bagaglio et al., 2017; Irwin et al., 2016). Due to the low fidelity of viral RdRps in general, the mutation frequency is estimated to be 10^−4^ to 10^−6^ errors per nucleotide (Lauring et al., 2013). The high mutation frequency and high replication rate of viral RNA copies enable the viruses quickly to adapt to changes in the environment, including the introduction of antiviral drugs. Identification of mutations conferring antiviral resistance provides information not only about the risk of generation of drug-resistant mutants but also helps to elucidate molecular mechanisms of the antiviral action. This is an integral and essential part of development and testing of any new antiviral drug (Irwin et al., 2016).

In the present study, we identified and described a specific amino acid substitution in the TBEV NS5 polymerase that confers resistance to Galidesivir. This substitution had only a limited effect on viral reproduction *in vitro*, but had a cost on viral fitness when tested *in vivo*, using mice. We also used structural modeling to link Galidesivir resistance to a molecular change in the NS5 RdRP active site that affects nucleotide incorporation. Our findings are important for understanding the mechanism of action of Galidesivir and for the use of this molecule as an antiviral drug against TBEV and other emerging RNA viruses. In addition, we highlight the discovery of a potential contribution of viral RdRp to flavivirus neurovirulence.

## Results

### TBEV resistant to Galidesivir has two amino acid substitutions in the NS5 protein

Studies of drug-resistant virus mutants are crucial for understanding molecular interactions of antiviral drugs with target viral proteins, as well as for development of efficient and specific antiviral therapies. In order to select TBEV resistant to Galidesivir, the virus was serially passaged in PS cell monolayers in the presence of increasing concentrations of Galidesivir up to 50 μM (Figure 1B); this process resulted in selection of two independent drug-resistant TBEV mutant strains. Whole genome sequencing of the passaged viruses revealed, that both selected TBEV mutants carried a single amino acid change E460D, which corresponds to the nucleotide substitution G9045T in the NS5 gene (Figure 1C,D). Sequencing of viruses after each passage showed, that this mutation was acquired after 5 passages. In passages 6 and 7, mixed mutated and wild-type genotypes were detected; from passage 8 onwards, the mutated genotype dominated until the end of the experiment (passage 9) (Figure 1D). Interestingly, in one of the selected TBEV mutants, an additional amino acid change Y453H was detected; this mutation, which was acquired after 8 passages, corresponds to the nucleotide substitution T9022C in the NS5 gene (Figure 1D). Mutations E460D and Y453H were not present in the wild-type virus passaged in the absence of the selection agents. Both mutations mapped to the active site of the RdRp domain of the NS5 protein. The in vitro selected TBEV mutants (denoted as E460D and E460D/Y453H) were further evaluated for their sensitivity/resistance to Galidesivir at concentrations ranging from 0 to 50 μM and compared with the mock-selected wild-type virus (Figure 2A). Whereas *in vitro* replication of wild-type was completely inhibited by Galidesivir at a concentration of 12.5 μM (EC_50_ value of 0.95 ± 0.04 μM), both mutants were approximately 7-fold less sensitive to the compound, showing EC_50_ values of 6.66 ± 0.04 and 7.20 ± 0.09 μM for E460D and E460D/Y453H, respectively (Figure 2A, Table 2).

**Fig 2.**
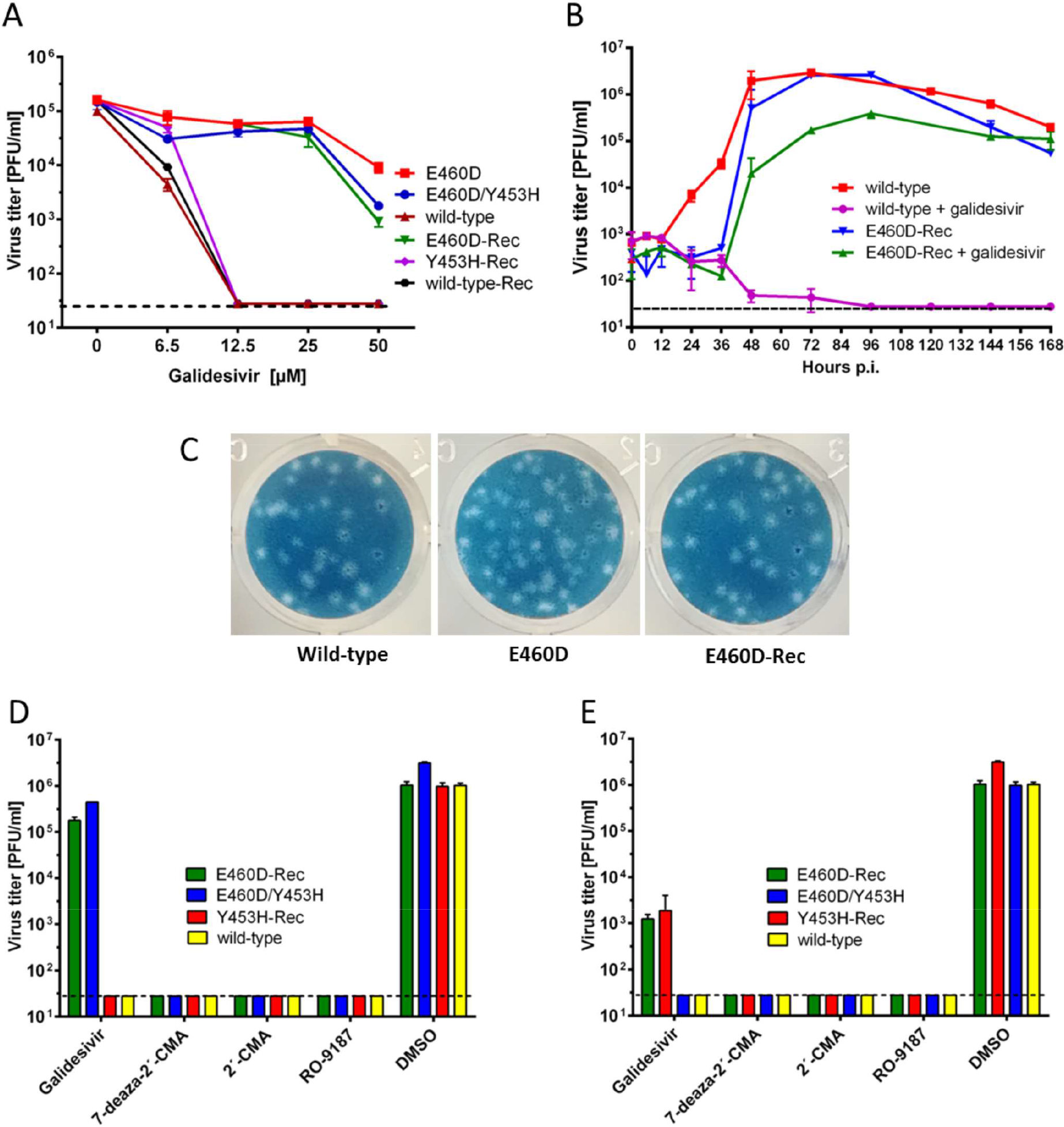
Phenotypic properties of TBEV mutant resistant to Galidesivir *in vitro.* (A) The dose-response curves for TBEV mutants E460D, E460D/E453H, E460D-Rec, Y453H-Rec, and corresponding wild-types grown in PS cells in the presence of Galidesivir at indicated compound concentrations. Only TBEVs bearing the E460D mutation (i.e., E460D, E460D/E453H, and E460D-Rec) were resistant to Galidesivir, indicating that this mutation is responsible for the resistance phenotype. (B) Growth kinetics of the E460D-Rec mutant and wild-type TBEV in the presence (25 μM) or absence (0 μM) of Galidesivir within the 7-day experimental period to assess the replication efficacy of the mutant TBEV in PS cells. (C) Plaque morphology of the E460D and E460D-Rec mutants was assessed in PS cell monolayers and compared with the wild-type virus. (D-E) The sensitivity/resistance profiles of the E460D/Y453H, E460D-Rec, and Y453H-Rec to diverse nucleoside inhibitors (concentrations of 25 μM (D) and 50 μM (E)) were evaluated in PS cells and compared with the corresponding wild-type TBEV. The mean titers from two independent experiments, each performed in triplicate are shown and error bars indicate standard errors of the mean. The horizontal dashed line indicates the minimum detectable threshold of 1.44 log_10_ PFU/mL.

**Table 1.**
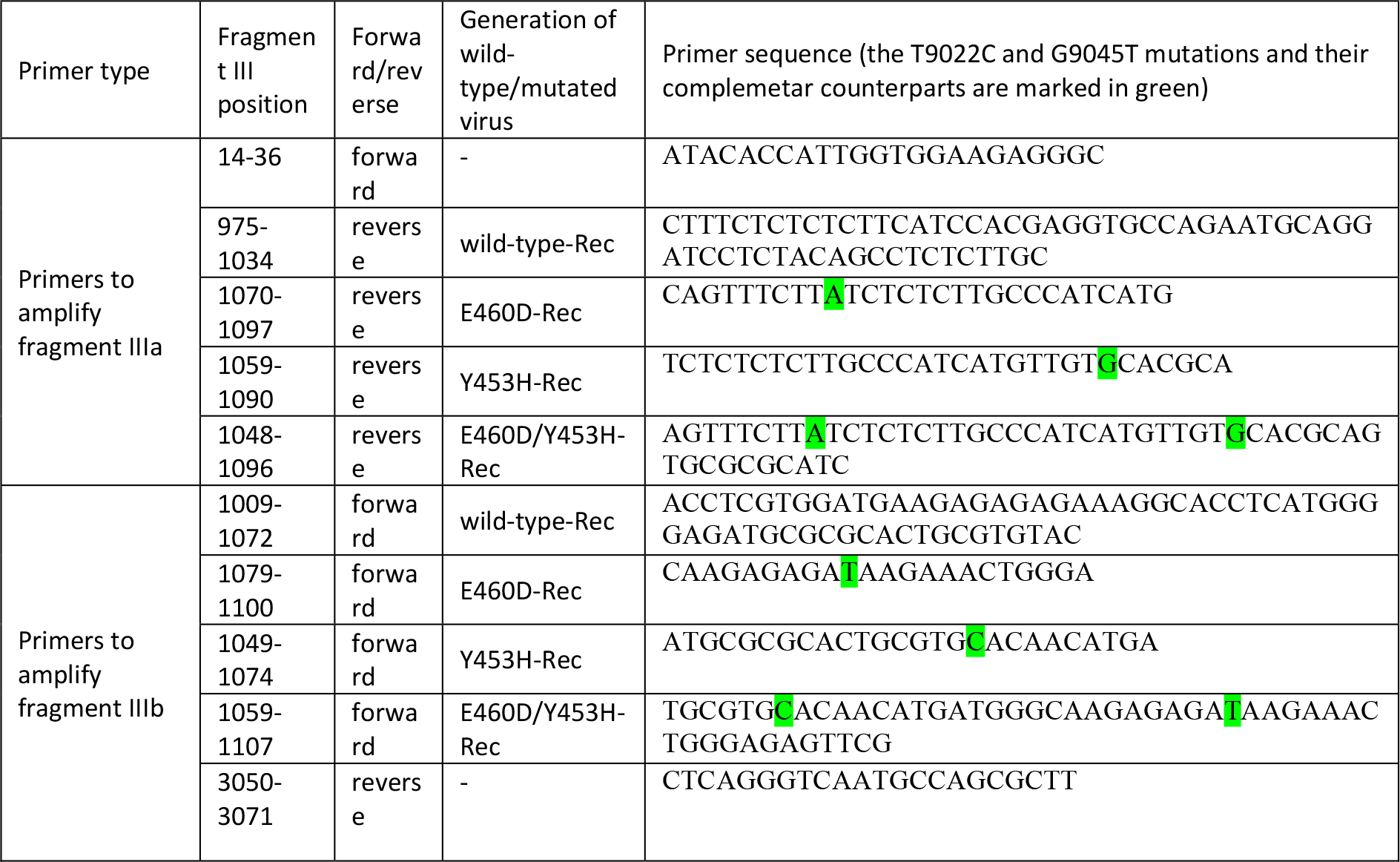
Unmodified and mutated primers used to generate two overlapping amplicons of fragment III (fragment IIIa and IIIb), i.e. to produce recombinant wild-type and mutated viruses

**Table 2.** Inhibitory properties of Galidesivir for the obtained TBEVs

| Virus | Method to generate wild-type/mutated virus | EC<sub>50</sub> [μM]* (fold increase compared to wild-type value) |
| --- | --- | --- |
| E460D | *In vitro* selection<br>(passaging in PS cells) | 6.66 ± 0.04 (7.01) |
| E460D/ Y453H |  | 7.20 ± 0.09 (7.57) |
| wild type |  | 0.95 ± 0.04 |
| E460D-Rec | Reverse genetics | 6.32 ± 1.01 (7.90) |
| Y453H-Rec |  | 0.81 ± 0.01 (1.01) |
| E460D/ Y453H-Rec |  | not rescued |
| wild-type-Rec |  | 0.80 ± 0.01 |

\*Determined from three independent experiments. Expressed as a 50% reduction of viral titers and calculated according to the Reed-Muench method. PS cells were infected with the virus (MOI of 0.1) and cultivated at Galidesivir concentrations ranging from 0 to 50 μM for three days.

### Site-directed mutagenesis confirms that E460D determines TBEV drug resistance

In order to demonstrate the direct effect of the amino acid substitutions E460D and Y453H on TBEV phenotype, the appropriate mutations were introduced into recombinant TBEV strains generated by the rapid reverse genetic approach based on the use of subgenomic overlapping DNA fragments (Aubry et al., 2014; Driouich et al, 2018). The entire TBEV strain Hypr genome flanked at the 5’and 3’untranslated regions by the pCMV and HDR/SV40pA was *de novo* synthesized in three double stranded DNA fragments of approximately 4.4, 4.5 and 3.1 kb in length, overlapping by 80 to 120 pb. The substitutions E460D (G9045T) and Y453H (T9022C) were introduced into the NS5 gene located on fragment III using mutagenic PCR primers (Figure 4). After transfection of the sub-genomic fragments into permissive BHK-21 cells, the following recombinant TBEV strains were successfully rescued: E460D-Rec (E460D substitution in the NS5 gene), Y453H-Rec (Y453H substitution in the NS5 gene), and recombinant wild-type (no introduced mutations). The presence of the E460D and Y453H substitutions in the viral genomes was confirmed by whole-genome sequencing of all recombinant viruses. Despite repeated attempts, the E460D/Y453H-Rec mutant (both mutations E460D and Y453H in the NS5 gene) was not rescued from the transfected BHK-21 cell culture. E460D-Rec was found to be 7.9-fold less sensitive to Galidesivir than engineered wild-type, showing an EC_50_ value of 6.32 ± 1.01 μM (Table 2). On the other hand, Y453H-Rec was highly sensitive to Galidesivir; the replication of this mutant strain was completely inhibited at 12.5 μM, showing an EC_50_ value of 0.81 ± 0.01 μM (Table 2). Thus, the results indicate, that the E460D (not Y453H) substitution is solely responsible for the drug-resistant phenotype of TBEV (Figure 2A).

### The E460D mutation has only a moderate effect on TBEV replication in cell culture

To characterize the phenotypic properties of the drug-resistant mutant *in vitro*, growth kinetics (Figure 2B) and plaque morphology (Figure 2C) of the recombinant TBEV mutant E460D-Rec were assayed in cultures of PS cells and compared with the wild-type virus. The wild-type virus amplified in the absence of Galidesivir (Figure 2B, red line), showed a short lag-period within intervals 0 – 12 hours p.i. Starting 24 hours p.i., the wild-type TBEV exerted an exponential increase in virus infectivity reaching a peak titre of 3×l0^6^ PFU/mL within 72 hours p.i. and gradually declining thereafter. In contrast, the presence of Galidesivir (25 μM) completely inhibited replication of the wild type virus (Figure 2B, violet line).

In comparison with the wild-type virus, the E460D-Rec mutant cultured in the absence of Galidesivir (Figure 2B, blue line) showed a prolonged lag-period within intervals 0 – 36 hours p.i. However, subsequently, the infectivity of the mutant virus increased exponentially reaching a peak of 2.6×l0^6^ PFU/mL at 72 hours p.i. After that, the titre gradually declined to 5.5×l0^4^ PFU/mL. The considerably longer lag-period of the E460D-Rec mutant could be explained by a slightly decreased replication capacity (attenuation) of the mutant, when amplified in PS cell culture. The decrease in replication capacity of E460D-Rec was manifested particularly in the first few hours after cell culture infection and was no longer detecable in the later stages of the infection.

The E460D-Rec mutant cultured in the prensence of Galidesivir (25 μM) (Figure 2B, green line) also exerted an extended lag-period within intervals 0-36 hours p.i.; there was even a moderate decrease in viral titers after 36 hours p.i. Starting 48 hours p.i., the mutant showed an exponential infectivity and reached a peak titre of 3.9×l0^5^ PFU/mL at day 96 hours p.i. The results clearly indicate, that the resistance of TBEV to Galidesivir is only partial; the growth of the mutant strain in the presence of Galidesivir was partially inhibited compared to wild-type (Figure 2B, red line) or the E460D-Rec mutant grown in the absence of Galidesivir (Figure 2B, blue line).

The plaque morphology of the drug-resistant TBEV mutant was similar to that of wild-type virus; both viruses produced large and clear plaques which were round and regular in shape and did not change in shape and size during all the consecutive passages (Figure 2C). The similarity in plaque morphology of drug-resistant and wild-type TBEVs is in agreement with similar growth kinetics of both viruses and supports our assumption that the mutation E460D affects the viral replication in PS cell culture only to a limited extent.

### The E460D TBEV mutant is sensitive to 7-deaza-2’-C-methyladenosine, 2-C-methyladenosine and 4’-azido-aracytidine

To test whether or not E460D and Y453H substitutions affect sensitivity of TBEV to structurally different nucleoside inhibitors we evaluated selected nucleoside analogues with previously reported anti-TBEV activity (Eyer et al., 2016), for their capacity to inhibit *in vitro* replication of Galidesivir-resistant TBEV. Inhibitory effects of 7-deaza-2’-*C*-methyladenosine, 2’-*C*-methyladenosine, and 4’-azido-aracytidine (RO-9187) at concentrations of 25 and 50 μM were not affected in the E460D/Y453H mutant which was obtained by serial sub-culture in PS cells in the presence of Galidesivir (EC_50_ >50 μM) (Figure 2D,E). The same drug-sensitivity profile, characterized by complete inhibition of virus replication, was determined for two recombinant TBEVs generated by reverse genetics, E460D-Rec and Y453H-Rec (EC_50_ >50 μM). Wild-type virus was used as a positive control in this *in vitro* antiviral study (Figure 2D,E).

### Mouse neuroinvasiveness of the E460D TBEV mutant is highly attenuated

The degree of neuroinvasiveness of the E460D TBEV mutant was assessed in BALB/c mice and was compared with that of wild-type virus. Adult BALB/c mice were infected subcutaneously with 10^3^ PFU of either virus and survival rates and clinical signs of neuroinfection were monitored for 28 days. Wild-type virus produced fatal infections in all mice, with mean survival times of 11 ± 2.2 days; infected mice showed severe signs of disease, such as ruffled fur, hunched posture, tremor and hind leg paralysis (Figure 3A,C). In contrast, all mice infected with the drug-resistant TBEV mutant E460D (obtained by serial *in vitro* sub-culture in the presence of Galidesivir) survived (*p*<0.0001), displaying no clinical signs of TBEV infection through the entire 28-day experimental period (Figure 3A,C). The same survival data (100% survival rate, *p*<0.0001) and clinical scores (no signs of neuroinfection) were obtained, when recombinant TBEV mutant (E460D-Rec) was used for mouse infection using the same infectious dose and administration route (Figure 3B,D).

**Fig 3.**
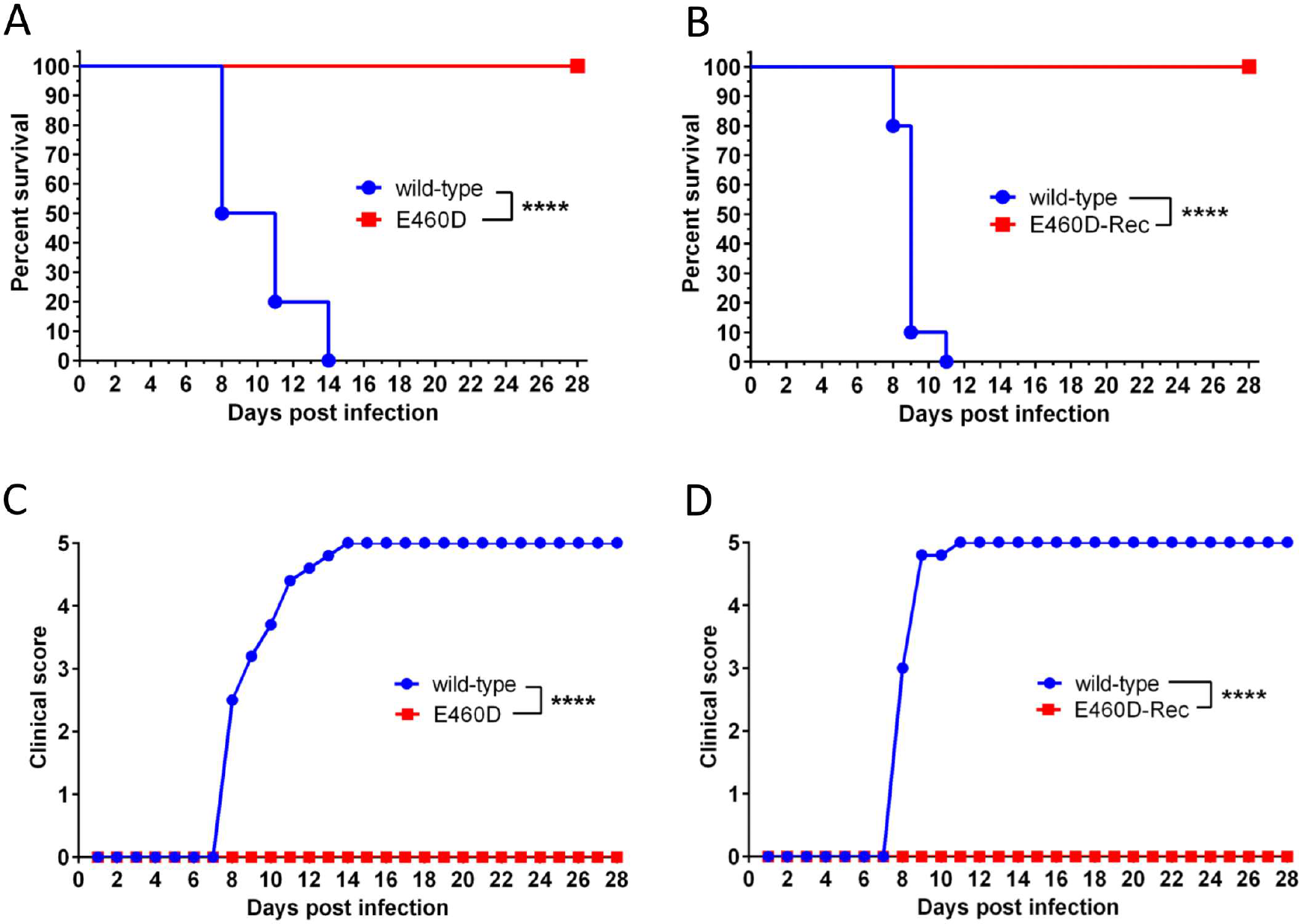
Phenotypic properties of the TBEV mutant resistant to Galidesivir in mice. The extent of neuroinvasiveness of the E460D and E460D-Rec TBEV mutants was investigated in BALB/c mice and compared with that of the wild-type virus. Adult BALB/c mice were infected subcutaneously with 10^3^ PFU of either virus and survival (A-B) and clinical scores of the neuroinfection (C-D) were monitored for 28 days. ****, *p*<0.0001.

### Rapid re-induction of wild-type genotype in the absence of Galidesivir

In general, mutant genotypes rapidly revert to the wild-type in the absence of the selection agents. Indeed, after three passages in PS cells in the absence of Galidesivir a pool comprising mutated and wild-type genotypes was detected. Moreover from passage 5 onwards, the wild-type genotype dominated in the infected PS cell culture (Figure 1E). Interestingly, the codon GAT encoding an aspartic acid in the E460D mutant had changed to the codon GAA in the revertant, not to GAG as seen in mock-selected wild-type virus; both codons, GAA (in the revertant) and GAG (in the wild-type), are synonymous and encode a glutamic acid residue (Figure 1F). In contrast, after 6 passages in the presence of Galidesivir, at concentrations up to 25 μM, the E460D substitution was retained; even at low concentrations of Galidesivir (6.25 μM) and did not result in reversion to the wild-type genotype (data not shown).

### Galidesivir migration towards the TBEV RdRp active site

The deduced position of Galidesivir bound within the TBEV RdRp active site is shown in Figure 4A with its phosphate tail forming close contacts with the manganese cofactors. The E460D (and Y453H) mutation is located at motif F of the flavivirus RdRp finger domain. The flavivirus RdRp motif F is highly flexible and occludes the NTP-tunnel in the nucleotide bound RdRp structure (Valdés et al., 2016). Therefore, the apo-structure was used for stochastic simulations to compare and contrast how Galidesivir approaches the TBEV active site in the wild type and mutant RdRps. At the active site of the wild type TBEV RdRp, the ribose of Galidesivir is at a 5.7 Å distance from amino acid (aa) position 460. The missing carbon in Asp increases this distance (7 Å) compared to Glu. This increased distance, caused by the mutation, results in more steric freedom for Galidesivir as it approaches the active site (Figure 4B,C).

**Fig 4.**
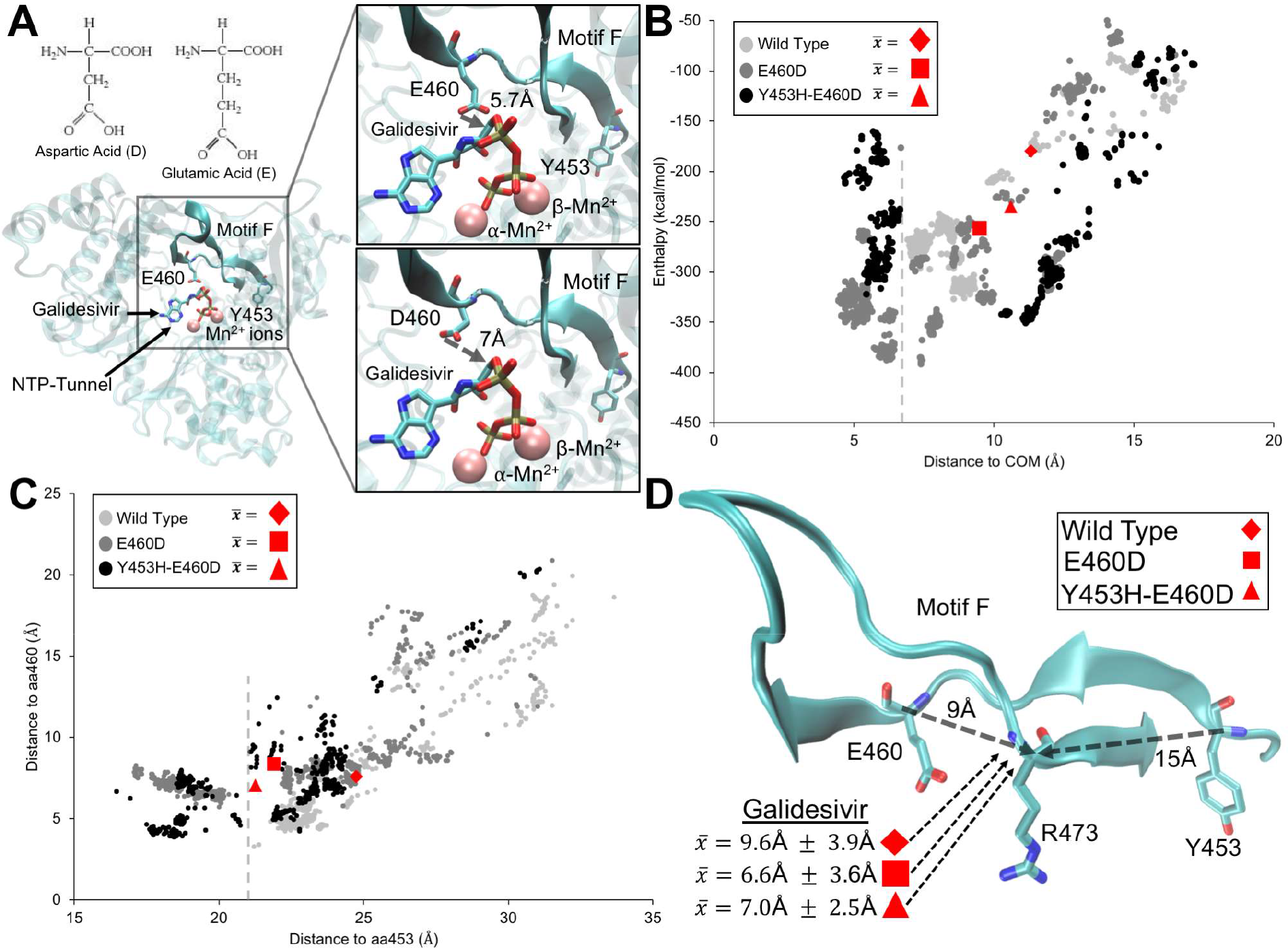
Proposed positioning of Galidesivir and its migration in the wild type and mutant TBEV polymerase. (A) The TBEV polymerase structure shown at 180° from its canonical right-hand conformation. Motif F and the residues subject to mutation are highlighted with Galidesivir and two Mn^2+^ ions (box). The NTP-tunnel is indicated by the arrow. The insets show the distance from the oxygen of E460 to the oxygen 5’ of Galidesivir and the distance to the mutant, D460 - shown at a 90° orientation. The Lewis structures show the missing carbon in Asp compared to Glu. (B) The x-y scatter plot is the Galidesivir binding enthalpy in kcal/mol (y-axis) during its migration to the hypothetical center of mass (COM) position within the TBEV polymerase active site (x-axis). The averages of the three replicates are shown as red geometric shapes (legend). (C) The x-y scatter plot is the Galidesivir distance to amino acid (aa) residue positions 453 (x-axis) and 460 (y-axis) that are subject to mutation in the TBEV polymerase. (D) Structural representation of motif F in a similar orientation as in (A) showing the average distance of Galidesivir, as red geometric shapes, to the interrogating residue R473. The distances of Y453 and E460 to the interrogating residue R473 (carbon backbone) are indicated by arrows.

There is a ∼6.7 Å distance cutoff by Galidesivir approaching the wild type TBEV RdRp active site that is explored closer (<5 Å) in both mutant types (Figure 4B). The single RdRp mutation, E460D, however, has lower enthalpic ligand binding values compared with the double mutant, E460D/Y453H. These lower enthalpic values indicate a favorable ligand binding and conformation. The distinct differences in Galidesivir exploration between the wild type and mutant RdRps are shown by the average COM distance/enthalpy in Figure 4B, specifically those by the single E460D mutation (wild type: 11.3 Å ± 4.6 Å /-179.9 kcal/mol ± kcal/mol, E460D: 9.5 Å ± 4.9 Å /-257.3 kcal/mol ± 121.2 kcal/mol; E460D/Y453H: 10.6 Å ± 4.5 Å /-235 kcal/mol ± 96.2 kcal/mol). The exploration difference between the RdRps is also demonstrated by the distance of Galidesivir to aa position 453, indicated by the cutoff distance ∼21 Å in the wild type RdRp (Figure 4C). Average distances to aa453 are, wild type 24.7 Å ± 3.1 Å/7.6 Å ± 4.2 Å, E460D 21.9 Å ± 3.5 Å/8.4 Å ± 2.7 Å and E460D/Y453H 21.3 Å ± Å /7.1 Å ± 2.7 Å. The Galidesivir distances to aa460 is maintained at ∼7.7 Å in all three RdRp types on average.

The Arg interrogation residue of motif F is highly conserved in flavivirus RdRps and its electrostatic interaction facilitates incorporation of favorable/correct incoming nucleotides (Valdés et al., 2016; Butcher et al., 2001; Bressanelli et al., 2002). Figure 4D shows that the distances of both mutation sites to Arg473 of the TBEV RdRp within motif F are located 9 Å and 15 Å away. The average distances of Galidesivir to Arg473 during the stochastic simulations are also noted. Due to the steric freedom within the active site caused by E460D (Figure 4A), both substitutions permit closer Galidesivir interactions with the interrogation Arg473. Close proximity to the interrogation residue will thereby increase electrostatic interactions with Galidesivir.

## Discussion

Galidesivir is an adenosine analogue originally developed for filovirus infection treatment (Ebola and Marburg) with high antiviral potency against a broad spectrum of RNA viruses (Warren et al., 2014; Taylor et al., 2016), including TBEV (Eyer et al., 2017b) and other medically important arthropod-borne flaviviruses (Warren et al., 2014; Julander et al., 2014; Eyer et al., 2017b; Julander et al., 2017). Currently, this compound entered first-in-human clinical studies that focused on intramuscular administration in healthy volunteers showing promising pharmacokinetics properties and good tolerability (Taylor et al., 2016). This makes Galidesivir a promising candidate drug to treat patients with TBE or with other flaviviral infections. However, antiviral therapy based on small molecule inhibitors of viral replication can be accompanied by rapid evolution of drug-resistance which can abolish the progress of infection treatment and finally lead to the failure of the therapy. Therefore, for each new antiviral agent, the risks of resistance are important to assess in terms of (i) identification of key mutations conferring virus drug-resistance and (ii) phenotype characterization of drug-resistant mutants.

Serial *in vitro* passaging of TBEV in the presence of increasing concentrations of Galidesivir (up to 50 μM) resulted in generation of two drug-resistant TBEV mutants which were approximately 7-fold less sensitive to Galidesivir than the mock-selected wild-type virus. The first TBEV mutant was characterized by a single amino acid change E460D; the other one carried two amino acid changes, E460D and Y453H. Both mutations mapped to the active site of the viral RdRp. Location of the resistance-associated mutations within the viral RdRp active site is essential to understand the mechanism of action of Galidesivir; this compound prevents the binding of subsequent nucleotides to the RdRp active site, being considered a non-obligate chain terminator of viral RNA synthesis (Warren et al., 2014; De Clercq and Neyts, 2009). Single amino acid changes within the RdRp were previously identified in flaviviruses resistant to structurally different nucleoside analogues, as exemplified by the mutations S603T, S604T and S282T conferring a high-level resistance to 2’-*C*-methylated nucleosides in TBEV, Zika virus and hepatits C virus, respectively (Eyer et al., 2017a; Hercik et al., 2017; Migliaccio et al., 2003). In Alkhurma haemorrhagic fever virus, the mutation S603T was associated with additional amino acid substitutions located in the NS5 RdRp active site, particularly with C666S and M644V (Flint et al., 2014).

Using a previously described reverse genetics system (Aubry et al., 2014; Driouich et al., 2018) we have demonstrated, that the E460D mutation alone is crucial for resistance of TBEV to Galidesivir; the recombinant E460D-Rec mutant was approximately 7-fold less sensitive to Galidesivir compared with wild-type. On the other hand, the growth kinetics of the Y453H-Rec mutant was almost indistinguishable from that of wild-type virus; it is likely that Y453H can represent a compensation mutation or was acquired randomly. Because of the unique structural features of Galidesivir (C-glycosidic bond and furanose oxygen on the ribose ring replaced by nitrogen) (De Clercq, 2016), no cross-resistance was seen to structurally different nucleoside analogues, such as 7-deaza-2’-*C*-methyladenosine, *2’-*C*-*methyladenosine and 4’-azido-aracytidine.

The E460D TBEV mutant showed similar growth kinetics to the wild-type virus, when cultured *in vitro* on PS cell monolayers. Although, a decreased replication capacity of the E460D mutant was observed in the first few hours after cell culture infection (0 – 12 hours p.i.), both viruses reached a peak titre of about 10^5^ – 10^6^ PFU/mL at days 2-4 after infection. Similarly, the plaque morphology of the E460D mutant and wild-type virus were almost identical to each other; large, clear and round plaques reflected rapid and aggressive spread of both mutant and wild-type viruses in PS cell cultures. Thus, the E460D TBEV mutant differs from the recently isolated S603T TBEV mutant resistant to 2’-*C*-methyladed nucleosides; the S603T mutant exerted significantly decreased replication capacity in PS cells and completely different plaque morphology (small, turbid plaques) compared with the wild-type virus (Eyer et al., 2017a). Our results demonstrate that antiviral resistance developed against two structurally different nucleoside analogues having the same mechanism of action can result in different effects on viral replication capacity in cell culture. Interestingly, in some drug-resistant virus mutants the cell-type dependent replication fitness was observed, as seen in chikungunya virus resistant to T-705 showing the attenuated phenotype in mosquito cell culture, whereas the replication fitness in Vero cells was similar to that of the wild type (Delang et al., 2018).

Although, the introduction of the E460D mutation affects amplification of the virus in PS cell culture (i.e., *in vitro*) only slightly, the E460D substitution resulted in a total loss of neuroinvasiveness for mice, *in vivo;* the E460D-infected animals all survived and displayed no clinical signs of neuroinfection during the 28-day experimental period. In contrast, infection with wild-type virus resulted in fatal infections for all animals. We propose that the E460D substitution could affect viral capacity to cross host barriers or responses that restrict the virus infection and translocation to the target tissues/organs *in vivo* but such possibilities do not occur during virus replication in PS cell culture, i.e. *in vitro.* Nevertheless, similar levels of attenuation *in vivo* have previously been reported for drug-resistant RNA or DNA viral mutants, i.e. for TBEV resistance to 2’-*C*-methylated nucleosides (Eyer et al., 2017a), chikungunya virus resistance to T-705 (Delang et al., 2018), Ribavirin-resistance to porcine reproductive and respiratory syndrome virus (Khatun et al., 2016), vaccinia virus resistance to acyclic nucleoside phosphonates (Gammon et al., 2008), and pleconaril-resistance to coxsackievirus (Groarke and Pevear, 1999).

The mutant genotype rapidly reverted to wild-type, when the virus was cultured in PS cell monolayers in the absence of selection agents. However, reversion was not observed, when the virus was cultured in the presence of Galidesivir in concentrations ranging from 6.25 to 25 μM. Thus, under the selection pressure of Galidesivir, the mutation provides a replicative advantage over wild-type variants in the virus quasispecies population resulting in predominance of the mutant in the infected cell culture, despite the fact that the replication characteristics of both variants in cell culture are similar.

We conclude that the resistance of TBEV to the nucleoside analogue Galidesivir is conferred by the single amino acid substitution E460D in the NS5 protein. Although this subtle mutation in the active site of the viral RdRp occurs after a few *in vitro* passages of TBEV in the presence of Galidesivir, the E460D TBEV mutant displays dramatically attenuated phenotype in mice showing high survival rates and reduction of clinical signs of neuroinfection. The stochastic molecular simulations indicate that the steric freedom caused by the E460D mutation increases close electrostatic interactions between Galidesivir and the interrogation residue of the TBEV RdRp motif F. Such close electrostatic interactions will reject the analogue as an incorrect nucleotide. The E460D substitution did not confer cross-resistance to unrelated antiviral nucleoside analogues, such as 7-deaza-2’-*C*-methyladenosine, 2’-*C*-methyladenosine and 4’-azido-aracytidine. This suggests that a combination treatment based on two or more inhibitors could be a possible strategy in order to minimize the risk of the emergence of viral drug resistance following therapeutic treatment with Galidesivir.

## Materials and methods

### Ethics statement

This study was carried out in strict accordance with the Czech national law and guidelines on the use of experimental animals and protection of animals against cruelty (the Animal Welfare Act Number 246/1992 Coll.). The protocol was approved by the Committee on the Ethics of Animal Experiments of the Institute of Parasitology and of the Departmental Expert Committee for the Approval of Projects of Experiments on Animals of the Academy of Sciences of the Czech Republic (Permit Number: 29/2016).

### Virus, cells, and antiviral compounds

A well-characterized, low-passage TBEV strain Hypr (Pospisil et al., 1954), a member of the European TBEV subtype, was used in this study. Before use, the virus was sub-cultured intracerebrally six times in suckling mice. Porcine kidney stable (PS) cells (Kozuch and Mayer, 1975) were used for viral subculture, selection of drug-resistant viruses, viral growth kinetics studies, and plaque assays. The cells were cultured at 37 °C in Leibovitz (L-15) medium supplemented with 3% newborn calf serum and a 1% mixture of Penicillin and glutamine (Sigma-Aldrich, Prague, Czech Republic). BHK-21 cells (obtained from the American Type Culture Collection [ATCC]), used for transfection of Hypr-derived subgenomic fragments, were cultured at 37 °C with 5% CO2 in Minimal Essential Medium (MEM) containing 7% bovine serum, 1% Penicillin/Streptomycin and glutamine. Galidesivir (BCX4430) and 4’-azido-aracytidine (RO-9187) were purchased from Medchemexpress (Stockholm, Sweden); 2’-*C*-methyladenosine and 7-deaza-2’-*C*-methyladenosine were from Carbosynth (Compton, UK). For *in vitro* studies, the test compounds were solubilized in 100% DMSO to yield 10 mM stock solutions.

### Selection of drug-resistant viruses

The *in vitro* selection of drug-resistant TBEV clones was performed, as described previously (Eyer at al., 2017a). Briefly, PS cells seeded in 96-well plates (2×l0^4^ cells per well) and incubated to form a confluent monolayer were infected with TBEV at a multiplicity of infection (MOI) of 0.1 and cultivated in the presence of 5 μM of Galidesivir. After 3 to 5 days, the culture medium was harvested and used for infection of fresh cell monolayers. Individual passages were performed with gradually increasing concentrations of Galidesivir as follows: passage 1 at 5 μM, passage 2 to 4 at 10 μM, passages 5 and 6 at 20 μM, passage 7 at 40 μM and passages 8 and 9 at 50 μM (Figure 2B). In parallel, control TBEV was also passaged in the absence of Galidesivir (with 0.5% (v/v) DMSO) as a mock-selected wild-type virus. After passage 9, the drug-resistant and control TBEVs were subjected to an additional subculture to prepare virus stocks for further testing (average titres were between 10^5^ – 10^6^ PFU/mL). The *in vitro* selection protocol was carried out in duplicate, resulting in two independent TBEV mutants, denoted as E460D and E490D/Y453H. In order to recover the revertant wild-type virus from the E460D population, the E460D virus pool was repeatedly sub-cultured in PS cells in the absence of Galidesivir; after 6 serial sub-cultures, the obtained revertant was amplified in PS cells to prepare a virus stock for further testing. Each of these viruses was subjected to full-length sequence analysis, sensitivity/resistance assessment to Galidesivir and other nucleoside analogues, and virulence characterization in mice.

### RNA isolation, PCR and whole-genome sequencing

RNA was isolated from growth media using QIAmp Viral RNA Mini Kit (Qiagen). Reverse transcription was performed using ProtoScript First Strand cDNA Kit (New England Biolabs) according to the manufacturer’s instructions for the synthesis of first strand cDNA, which was subsequently used for PCR amplification. To cover the whole genome of TBEV, 35 overlapping DNA fragments were produced by PCR as described previously Růžek et al., 2008). DNA was purified using High Pure PCR Product Purification Kit (Roche), according to the recommendations of the manufacturer. The PCR products were directly sequenced by commercial service (SEQme, Czech Republic) using the Sanger sequenation method. Both nucleotide and deduced amino acid sequences were analyzed using BioEdit Sequence Alignment Editor, version 7.2.0.

### Reverse genetics system for TBEV Hypr

Reverse genetics system used in this study was based on the generation of infectious subgenomic overlapping DNA fragments that encompass the entire viral genome as previously described (Aubry et al., 2014; Driouich et al., 2018). Three *de novo* synthesized DNA fragments cloned into a pUC57 vector were used in this study (GenScript, Piscataway, NJ, USA): fragment I (nucleotide position 1 to 3662), fragment II (nucleotide position 3545 to 8043), and fragment III (nucleotide position 7961 to 11100). The first and last fragment were flanked respectively in 5’and 3’with the human cytomegalovirus promoter (pCMV) and the hepatitis delta ribozyme followed by the simian virus 40 polyadenylation signal (HDR/SV40pA) (Figure 5).

**Fig 5.**
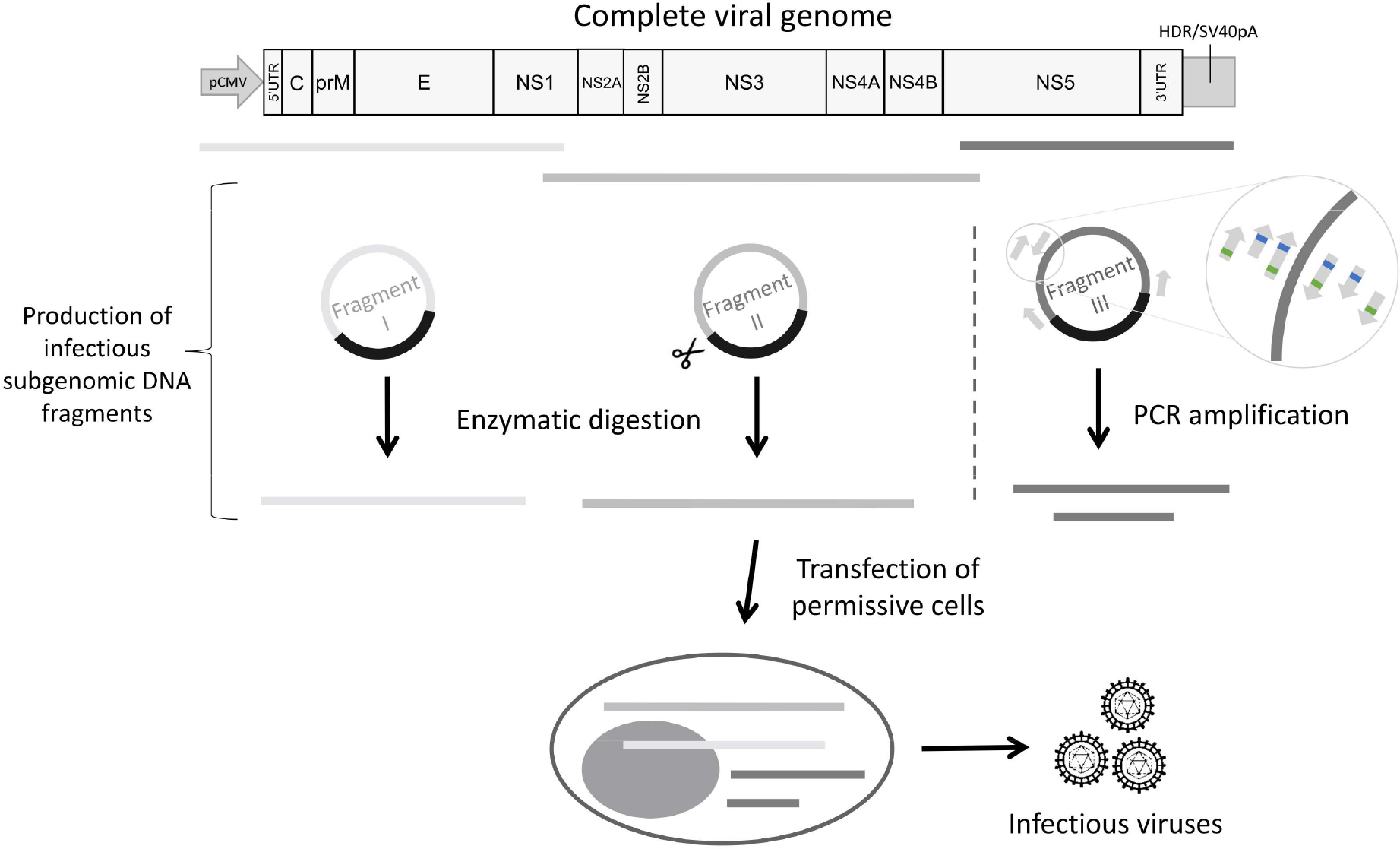
General overview of the reverse genetics method presented in this study. The reverse genetics method used in this study was based on the generation of infectious subgenomic overlapping DNA fragments that encompass the entire viral genome. Three *de novo* synthesized DNA fragments cloned into a pUC57 vector were used. The first and last fragment were flanked respectively in 5’and 3’with the human cytomegalovirus promoter (pCMV) and the hepatitis delta ribozyme followed by the simian virus 40 polyadenylation signal (HDR/SV40pA). Fragments I and II were generated using the SuPReMe method: plasmids were digested using restriction enzymes. Fragment III was used as template to generate by PCR two overlapping amplicons following the original ISA method. Unmodified primers were used to generate two unmodified amplicons *(i.e.* production of wild-type virus). Mutated primers located on the targeted region were used to generate two mutated amplicons (green and blue squares represent respectively mutations G9045T and T9022C). An equimolar mix of these four DNA fragments was used to transfect BHK-21 cells.

Fragments I and II were generated using the SuPReMe method (Driouich et al., 2018). Briefly, plasmids that contained the DNA fragments I and II were digested using respectively Agel/Fsel and Smal/Dral restriction enzymes (New England BioLabs, Ipswich, MA, USA) (Figure 4). Fragment III was used as template to generate by PCR two overlapping amplicons following the original ISA method (Aubry et al., 2014). Unmodified primers were used to generate two unmodified amplicons (*i.e.* production of wild-type virus). Mutated primers located on the targeted region were used to generate two mutated amplicons (i.e. production of mutated viruses, denoted as E460D-Rec, Y453H-Rec, and E460D/Y453H-Rec) (Table 1, Figure 4).

The PCR was performed using the Platinum SuperFI PCR Master Mix (Thermo Fisher Scientific, Prague, Czech Republic). The mixture (final volume, 50 μl) contained 45 μL of SuperMix, 2 μl of DNA template (fragment III) at 1 ng/μl. Assays were performed on a Biometra TProfessional Standard Gradient thermocycler with the following conditions: 94 °C for 2 min followed by 40 cycles of 94 °C for 15 s, 60 °C for 30 s, 68 °C for 5 min and a final elongation step of 68 °C for 10 min. Size of the PCR products was verified by gel electrophoresis and purified using an Amicon Ultra 0.5 ml kit (Millipore).

An equimolar mixture of these four DNA fragments was used for cell transfection. DNA-lipid complex was prepared as follows: 12 μl of Lipofectamine 3000 (Life Technologies) was diluted in 250 μL Opti-MEM medium (Life Technologies) and then mixed with a master solution of DNA which contained 3 μg of DNA and 6 zμl of P3000 reagent diluted in 250 μL Opti-MEM medium. After 45-min incubation at room temperature BHK-21 cells were transfected, as described previously (Aubry et al., 2015) and incubated for 5-7 days. Cell supernatant media were then harvested and serially passaged twice in fresh BHK-21 culture.

### Growth kinetics, dose-response studies and viral inhibition assays

To evaluate growth kinetics of drug-resistant TBEV mutants, PS cell monolayers incubated for 24 h in 96-well plates were treated with 200 μl of medium containing Galidesivir at concentrations of 25 μM (compound-treated cells) or 0.5% (v/v) DMSO (mock-treated cells) and simultaneously infected with TBEV; the MOI of 0.1 was used for all TBEV mutant/wild-type tested. The medium was collected from the wells daily at days 1 to 7 p.i. (three wells per interval); viral titres (expressed as PFU/mL) were determined by plaque assay as described previously (De Madrid and Porterfield, 1969; Eyer et al., 2015) and used to construct TBEV growth curves.

For dose-response studies, 200 μl of fresh medium containing Galidesivir at concentrations ranging from 0 to 50 μM was added to PS cell monolayers, infected with TBEV at an MOI of 0.1 and incubated for 3-4 days p.i. Then, the medium was collected from the wells and the viral titres were determined by plaque assay. The obtained viral titre values were used for the construction of TBEV dose-response/inhibition curves and for estimation of the 50% effective concentration (EC_50_) of the drug.

To measure the sensitivity/resistance of the obtained TBEV mutants to Galidesivir and several structurally unrelated nucleoside analogues in cell culture by viral titre inhibition assay, confluent PS cell monolayers cultured for 24 h at 37 °C in 96-well plates were treated with Galidesivir, 7-deaza-2’-*C*-methyladenosine, 2’-*C*-methyladenosine, or 4’-azido-aracytidine (RO-9187) at concentrations of 25 or 50 μM and simultaneously infected with TBEV at an MOI of 0.1 (3 wells per compound). As a mock-treated control, DMSO was added to virus- and mock-infected cells at a final concentration of 0.5% (v/v). The formation of cytopathic effect (CPE) was monitored visually using the Olympus BX-5 microscope to yield 70%-90% CPE in virus-infected cultures and viral titres were determined by plaque assays from cell culture supernatants.

### Mouse infections

To evaluate the virulence of the Hypr E460D mutant in mice, four groups of six-week old BALB/c female mice (purchased from AnLab, Prague, Czech Republic) were infected subcutaneously with TBEV (1,000 PFU/mouse) as follows: group 1 (n = 10), infected with *in vitro* mock-selected wild-type; group 2 (n = 10), infected with *in vitro* selected mutant (E460D); group 3 (n = 10), infected with recombinant wild-type TBEV; and group 4 (n = 10), infected with recombinant mutant (E460D-Rec). Survival rates of TBEV-infected mice were monitored daily over the 28-day experimental period. At the same time, controlling of illness symptoms and evaluation of clinical scores were performed in infected animals. Signs of sickness were evaluated as follows: 0 for no symptoms; 1 for ruffled fur; 2 for slowing of activity or hunched posture; 3 for asthenia or mild paralysis; 4 for lethargy, tremor, or complete paralysis of the limbs; 5 for death. All mice exhibiting disease consistent with clinical score 4 were terminated humanely (cervical dislocation) immediately upon detection.

### Stochastic molecular simulations

The unbound, apo-form of the TBEV RdRp used in this study is a homology-based predicted structure previously prepared for simulations in other published studies (Valdés et al., 2016, 2017). The stochastic molecular simulations were conducted using the online software, Protein Energy Landscape Exploration (PELE) that employs a Metropolis Monte Carlo algorithm which accepts or rejects a protein-ligand conformation based on its enthalpy. The PELE method and its applications are thoroughly explained online (https://pele.bsc.es/) and elsewhere (Madadkar-Sobhani et al., 2013; Borrelli et al., 2005) and therefore are briefly described here. The PELE software comprises three steps: a local protein and ligand perturbation, amino acid sidechain sampling, and a global minimization. These steps are repeated for a few thousand iterations resulting in a trajectory of a ligand (i.e., Galidesivir) approaching the active site of the target protein (i.e., TBEV RdRp). The first 25 snapshots were removed from the final trajectories prior to analysis since the stochastic simulations reached a relatively stable total enthalpy. Geometric and enthalpy measurements recoded via the PELE script were used for data analysis. The stochastic migration simulations were performed in triplicates, each with separate positioning of Galidesivir 20 Å ∼ 25 Å away from the centre of mass (COM) within the TBEV RdRp active site.

### Statistical analyses

Data are expressed as means ± SD, and the significance of differences between groups was evaluated using the Mann-Whitney *U* test or ANOVA. Survival rates were analysed by log-rank Mantel-Cox test. All tests were performed using GraphPad Prism 5.04 (GraphPad Software, Inc., San Diego, CA, USA). P-values < 0.05 were considered to be statistically significant.

## Acknowledgements

This study was supported by grant from the Ministry of Education, Youth and Sports of the Czech Republic (grant no. LTAUSA18016) (to LE), grant from the Ministry of Health of the Czech Republic (grant no. 16-34238A), and by Project “FIT” (Pharmacology, Immunotherapy, nanotoxicology; CZ.02.1.01/0.0/0.0/15_003/0000495) from the Ministry of Education, Youth and Sports of the Czech Republic, and Ministry of Agriculture of the Czech Republic (RO0518) (both to DR).

